# *Wolbachia* infection in *Aedes aegypti* does not affect its vectorial capacity for *Dirofilaria immitis*

**DOI:** 10.1101/2024.03.04.583272

**Authors:** Takahiro Shirozu, Maria Angenica F Regilme, Manabu Ote, Mizuki Sasaki, Akira Soga, Hiroki Bochimoto, Hidenobu Kawabata, Rika Umemiya-Shirafuji, Hirotaka Kanuka, Shinya Fukumoto

## Abstract

Mosquito-borne diseases such as dengue and filariasis are a growing public health concern in endemic countries. Biological approaches, such as the trans-infection of *Wolbachia pipientis* in mosquitoes, are an alternative vector control strategy, especially for arthropod-borne viruses such as dengue. In the present study, the effect of *Wolbachia* (wMel strain) on the vectorial capacity of *Aedes aegypti* for *Dirofilaria immitis* was studied. Our results showed that *Wolbachia* does not affect the phenotype of mosquito survival or the prevalence, number, and molting rate of third-stage larvae in both susceptible and resistant strains of *Ae*. *aegypti*. RNA-seq analysis of Malpighian tubules at 2 days post-infection with *D. immitis* showed the differentially expressed genes (DEGs) with and without wMel infection. No characteristic immune-related gene expression patterns were observed among the DEGs. No significant change in the amount of *Wolbachia* was observed in the *Ae. aegypti* after *D. immitis* infection. Our results suggest that infection of *D. immitis* in *Ae. aegypti* populations will not interfere with *Wolbachia*-based vector control strategies in dengue-endemic areas where cases of *D. immitis* are present. This study demonstrated the veterinary medical validity of a dengue control program using *Wolbachia*.

## Introduction

Mosquito-borne diseases such as dengue, malaria, and lymphatic filariasis are growing public health concerns, affecting 700,000 people worldwide [1, 2]. *Aedes* sp. is a potential mosquito vector for dengue, yellow fever, and chikungunya, all of which affect humans [1, 2]. Insecticide use is the most common method for mosquito-borne disease control [3]. However, the negative effects of insecticides on the environment and the development of insecticide-resistant mosquitoes have been reported previously [3]. Thus, biological control approaches, such as trans-infection of *Wolbachia pipientis* in the natural populations of the dengue vector *Aedes aegypti*, have recently been utilized as a better alternative to chemical-based approaches [4, 5]. The vectorial capacity of *Wolbachia*-infected mosquitoes for several viral infections have been reported, and *Wolbachia* strains wMel, wMelPop-CLA, and wAlbB have been found to show suppressive effects on the vectorial capacity for dengue, Zika, chikungunya, and West Nile viruses in *Ae*. *aegypti* [6–8].

*Wolbachia* is a maternally inherited intracellular bacterium found in insects and arthropods such as *Ae*. *aegypti* and *Culex pipiens* mosquito [9, 10]. *Wolbachia* can decrease the vectorial capacity for various pathogens, such as the dengue virus, in *Ae*. *aegypti* [6]. Moreover, *Wolbachia* spreads efficiently to the next generation because of its vertical transmission and cytoplasmic incompatibility [11]. *Wolbachia*-infected females produce *Wolbachia*-infected offspring regardless of whether the mating partner is infected or uninfected with *Wolbachia* [11]. When a *Wolbachia*-infected male mates with an uninfected female, some or all of the embryos die [11]. Successful establishment of *Wolbachia* (wMel strain)-infected mosquitoes in the field was observed in Cairns, Australia [12]. Several studies have also shown successful establishment of *Wolbachia* infections in natural *Aedes* populations in Brazil [13, 14], Australia, Vietnam [15], and New Caledonia [16]. Establishment of wMel-infected *Ae. aegypti* population has been reported to contribute to a 96% reduction in the incidence of dengue fever in Cairns [17]. Thus, *Wolbachia* can potentially be utilized to control mosquito-borne viral diseases, especially that caused by the dengue virus, in endemic areas.

However, to promote the use of *Wolbachia*, its effects on mosquito-borne diseases other than viral infections also require attention. *Wolbachia* also suppresses the vectorial capacity for parasitic infections transmitted by mosquitoes, with wMelPop and wAlbB showing suppressive effects on the vectorial capacity of *Anopheles gambiae* for *Plasmodium falciparum* [18] and that of *Anopheles stephensi* for *Plasmodium berghei* [19]. Conversely, *Wolbachia* has also been shown to increase the vectorial capacity of mosquitoes in some studies, as discussed below. wAlbB showed an adverse effect on the vectorial capacity for West Nile viruses in *Culex tarsalis* [20] and that for *P. berghei* in *An*. *gambiae* [21]. These studies suggest that *Wolbachia* infections in mosquitoes can positively or negatively affect the vectorial capacity for some viral and parasitic infections. Thus, further studies on the adverse effects of *Wolbachia* on pathogens other than dengue virus are essential for safe usage of *Wolbachia* in biological control approaches.

Filarial nematodes are parasites transmitted by mosquitoes such as *Aedes* and *Culex* [22]. Lymphatic filariasis, a human infection of public health importance that is caused by *Wuchereria bancrofti* and *Brugia malayi*, is a neglected tropical disease similar to dengue fever [23]. *Aedes* sp. is also a vector of *Brugia pahangi,* which causes lymphatic filariasis, and *Dirofilaria immitis,* which causes heartworm disease in canine, feline, and wild animals [22, 24]. *B. pahangi* is a useful experimental model for lymphatic filariasis [25, 26] and the risk of human zoonotic diseases [27]. *D*. *immitis* is also used as an experimental model for filarial transmission and development in mosquitoes [28, 29]. Dirofilariasis is endemic worldwide [24] and is recognized as a zoonosis [30]; therefore, it cannot be ignored from a public health perspective. Thus, the effect of *Wolbachia* on filarial parasites needs to be examined before expanding the use of *Wolbachia*-infected mosquitoes for dengue control.

The effect of *Wolbachia* infection on the vectorial capacity of mosquitoes for filarial infections has been reported in limited detail in the context of lymphatic filariasis, with wMelPop-infected *Ae. aegypti* [31] and wAlbB-infected *Ae. polynesiensis* [32] showing suppressive effects on *B. pahangi*. However, native wPolA-infected *Ae. polynesiensis* [32] and *Ae. pseudoscutellaris* infected with an unknown strain [33] did not show differences in vectorial capacity for *B*. *pahangi* in comparison with non-infected mosquitoes. In this regard, few reports have evaluated *D. immitis* even though it can cause lethal infections from a veterinary perspective. Mosquitoes infected with transgenic bacteria expressing a *Wolbachia* protein have shown suppressed vectorial capacity for *D. immitis* [34]. Therefore, *Wolbachia*-infected mosquitoes have the potential to be used for filariasis control.

In this study, we aimed to determine the effects of *Wolbachia* infection on the vectorial capacity of *Ae*. *aegypti* for *D. immitis* through a comparative analysis of the phenotypes of mosquitoes with and without *Wolbachia* (wMel). Specifically, the purpose of this study was to analyze the *D. immitis* infection phenotype, transcriptome, and amount of wMel in *Ae. aegypti* co-infected with wMel and *D. immitis*. Our results can help clarify whether *Wolbachia*-infected *Ae. aegypti* can be used as a new heartworm control strategy and if *Wolbachia* infection of *Ae. aegypti* is a safe method that does not increase the risk of spreading heartworm disease and shows no adverse effects.

## Methods

### Ethical approval

This study was conducted in strict accordance with the recommendations of the Guide for Laboratory Animals of the Obihiro University of Agriculture and Veterinary Medicine (OUAVM). The protocol was approved by the Committee of Animal Experiments of the OUAVM (permit number: 23-75 and 23-78). Mosquitoes infected with wMel were passaged using the blood of human volunteers (Ethics approval number: 201706-2 and 201801-2). Non-infected mosquitoes were passaged using the blood of mice or dogs.

### Experimental dogs and the *D. immitis* and *Ae. aegypti* strains

Female beagles (*Canis lupus familiaris*) aged 2–4 years were experimentally infected with the *D. immitis* SF1 strain, which was originally isolated in Japan [35], and reared under a veterinarian’s supervision. Six *Ae*. *aegypti* mosquito strains, MGYP2.tet, MGYP2, Liverpool (LVP)-Obihiro (OB), LVP-OB-wMel (OB-wMel), LVP-IB12 (IB12), and a cross between OB-wMel females and IB12 males (OB-wMel × IB12), were used in this study. MGYP2.tet and MGYP2 were provided by Monash University and Dr. Scott O’Neill, respectively. MGYP2 were artificially infected with *Wolbachia* wMel. MGYP2.tet was maintained free of wMel infection by tetracycline treatment. OB is an original *D. immitis*-susceptible strain identified in our laboratory [36]. OB-wMel was established by backcrossing female progenies of OB males and MGYP2 females with OB males showing maternal inheritance of *Wolbachia* to obtain a *D. immitis*-susceptible strain infected with wMel. Eight repeat backcross strains were confirmed to be infected with wMel by quantitative reverse transcription polymerase chain reaction (qRT-PCR). IB12 is a *D. immitis*-refractory strain obtained from BEI resources [37]. F1 progeny of OB-wMel × IB12 were obtained for analysis of the number of third-stage larvae (L3). The mosquitoes were maintained at 27°C and >80% humidity on a diet containing 5% fructose and a 12:12 h light:dark photoperiod.

### *D. immitis* infection

Approximately seven days after emergence, the mosquitoes were starved for 12 h before blood feeding. *D. immitis*-infected blood was collected from a *D. immitis*-infected dog kept in our laboratory and diluted with blood from a non-infected dog to adjust its concentration to 6-12 microfilaria (mf)/μL. The mosquitoes were artificially fed mf-infected blood via a plastic membrane (Parafilm; Bemis Flexible Packaging, WI, USA). Blood-fed mosquitoes until satiety were selected under CO_2_ anesthesia and kept in a cage for two weeks in an incubator under the same conditions as described above.

### Phenotypic analysis

The phenotypes of mosquito survival, prevalence rate, and the number of L3 in *D. immitis*-infected mosquitoes were analyzed. The number of surviving mosquitoes was recorded daily for 13 days post-infection (dpi). At 13 dpi, when most of the L3 had moved to the mosquito head, all surviving mosquitoes were dissected to investigate the prevalence rate and number of L3. Some of the L3 detected in each strain were also observed morphologically. The mosquito survival rate and the number of L3 were compared between each strain.

### L3 molting assay

L3 from OB and OB-wMel were subjected to the L3 molting assay *in vitro*. L3 on 13 dpi were collected in RPMI medium 1640 (Thermo Fisher Scientific, MA, USA) supplemented with penicillin, streptomycin, and 10% glucose. After washing twice with a medium containing 10% fetal bovine serum, L3 in good condition were selected based on their size and motility. Ten L3 per well were cultured in triplicate at 37°C and 5% CO_2_ for 1 week in 96-well plates for cell culture (Thermo Fisher Scientific) with 200 µL of medium containing 10% fetal bovine serum. Half of the medium was changed per day. The number of L3 that molted into L4 was recorded daily.

### qRT-PCR of wMel

The salivary gland (SG), midgut (MG), and Malpighian tubule (MT) (n > 10) were collected to confirm wMel infection in the backcross strain and to quantify the amount of wMel in mosquitoes after *D. immitis* infection. The time points for wMel quantification were day 0 (pre-blood feeding), 7, and 14 (non-blood feeding, non-infected blood feeding, and *D. immitis*-infected blood feeding). Genomic DNA was extracted from each sample using MagExtractor^TM^ Genome (Toyobo, Osaka, Japan). DNA samples were subjected to qPCR analysis to investigate the quantity of wMel relative to the homeobox protein homothorax (HTH) gene as mosquito DNA. The primer sequences used were wMel forward primer: TATTGAGCCTTCCTCGTACC, wMel reverse primer: TAGCATGCCGTTTTTCTGTA, HTH forward primer: TGGTCCTATATTGGCGAGCTA, and HTH reverse primer: TCGTTTTTGCAAGAAGGTCA [38]. The qPCR runs were performed using Applied Biosystems™ StepOne™ Real-Time PCR System (Thermo Fisher Scientific) and THUNDERBIRD^®^ SYBR qPCR Mix (Toyobo). Thermal cycling conditions were as follows: 95°C for 5 min, 40 cycles at 95°C for 15 s and 60°C for 45 s, followed by 95°C for 15 s, 60°C for 15 s, and 95°C for 15 s.

### RNA sequencing analysis

We compared the molecular basis of the host response in each mosquito strain after *D. immitis* infection by using RNA sequencing (RNA-seq) analysis. The mosquitoes (n > 10) were dissected at 2 dpi to collect MT tissues, and the pooled samples were stored at −80°C. RNA was extracted from the pooled MT samples using PureLink™ RNA Mini Kit (Thermo Fisher Scientific) and stored at −80°C. The RNA samples were subjected to RNA-seq using Illumina NovaSeq6000. RNA-seq analysis was conducted on the basis of read count data obtained by mapping raw read data to the reference genome data (*Aedes-aegypti*-LVP_AGWG_TRANSCRIPTS_AaegL5.2) from VectorBase (https://www.vectorbase.org/) using the CLC Genomics workbench software (CLC bio, Aarhus, Denmark). The log-intensity ratio (M) and log-intensity average (A) (MA) plot was created to detect differentially expressed genes (DEGs) between pairs of strains (MGYP2.tet vs. MGYP2, OB vs. OB-wMel, MGYP2.tet vs. OB, and MGYP2 vs. OB-wMel) based on the total read count data using R software (version 4.0.4). A heatmap was created to compare gene expression in each strain (MGYP2.tet, MGYP2, OB, and OB-wMel) based on Transcripts Per Million values using Heatmapper (http://www.heatmapper.ca) by selecting Average Linkage as the clustering method and Pearson as the distance measurement method. We focused on gene lists that showed upregulation by Toll activation [39], *Wolbachia* infection [40], and 48 h post-*Brugia malayi* infection [41]. Gene ontology analysis was conducted on the basis of the DEG list using the database for annotation, visualization, and integrated discovery (DAVID) as a bioinformatics resource.

### Statistical analysis

Statistical analyses were performed on the mosquito survival rate, number of L3, L3 molting rate, and relative amount of wMel DNA using GraphPad Prism 7 (GraphPad Software, CA, USA). Significant differences in mosquito survival rate and L3 molting rate were assessed using the Chi-square test for comparison between each strain, with and without wMel. The number of L3 was assessed by the Mann–Whitney test in a two-strain comparison (OB and OB-wMel) and by the Kruskal–Wallis test, a non-parametric one-way analysis of variance, in three-strain comparisons (OB, MGYP2, and MGYP2.tet and OB, IB12, and OB-wMel × IB12). The relative amounts of wMel DNA between strains and treatments were assessed using the Kruskal–Wallis test. Results with *p*-values of <0.05 were considered significant.

## Results

### Comparative analysis of phenotypes and L3 molting with and without wMel infection

As shown in Fig 1A-D, no significant differences were observed in the phenotypes of *D. immitis* infection between MGYP2.tet and MGYP2 mosquitoes. Mosquito survival and the number of L3 in MGYP2.tet and MGYP2 were significantly lower than those in the *D. immitis*-susceptible strain OB. The mosquito survival rate at 13 dpi was 19.4% (6/31) in MGYP2.tet, 26.9% (14/52) in MGYP2, and 89.4% (42/47) in OB (Fig 1A). The prevalence of L3 was 31.8% (7/22) in MGYP2.tet, 37% (10/27) in MGYP2, and 98% (96/98) in OB (Fig 1B). The number of L3 was 5.4 in MGYP2.tet, 8.3 in MGYP2, and 13.1 in OB (Fig 1C). L3 from MGYP2.tet and MGYP2 were morphologically normal in size and motility (Fig 1D).

**Fig 1.**
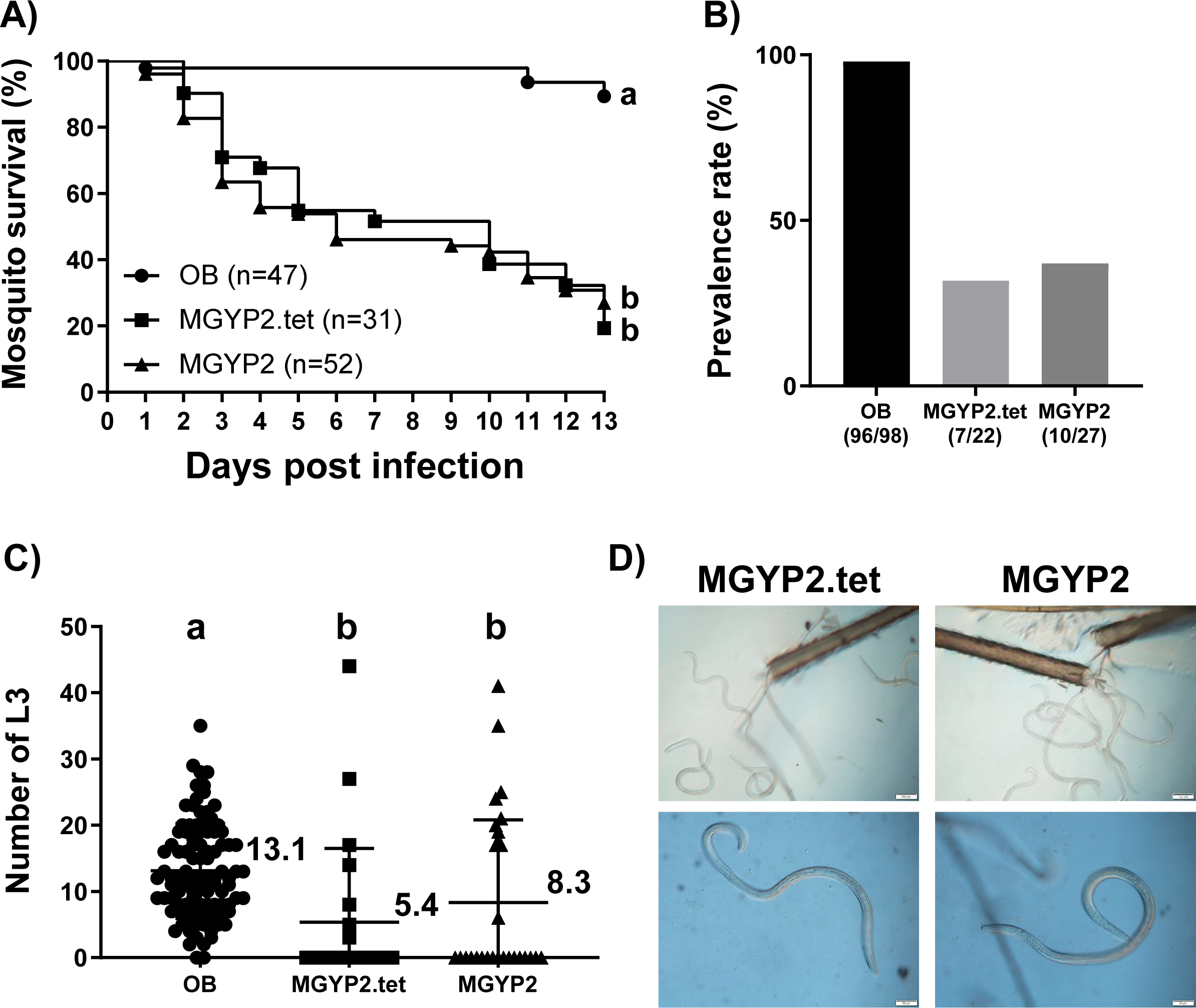
The *D. immitis* infection phenotype did not differ significantly between MGYP2 with and without wMel. *D. immitis* infection phenotype was compared between MGYP2.tet, MGYP2, and the susceptible OB as the positive control. The graphs show (A) mosquito survival up to 13 dpi, (B) prevalence rate of L3 on 13 dpi and (C) number of L3 per mosquito on 13 dpi in MGYP2.tet, MGYP2, and OB. The experimental sample size for mosquito survival and number of L3-positive/total as a measure of the prevalence rate are shown for each strain. The number of L3 is shown as the mean value with standard deviation (SD). Mosquito survival rate and number of L3 on 13 dpi were statistically analyzed. The data with different letters showed a significant difference (*p* < 0.05). The images indicate that (D) L3 from MGYP2.tet and MGYP2 on 13 dpi was normal. The upper and lower images show L3 in the proboscis of a mosquito and an enlarged image of L3, respectively. Scale bars of 100 or 50 μm are shown in the lower right corner of the image.

To evaluate the effect of wMel infection on the vectorial capacity for *D. immitis,* we established OB-wMel as a highly susceptible strain for *D. immitis* infection. In the qRT-PCR analyses, OB-wMel was confirmed to be infected with wMel, although no significant differences were observed (Fig 2A). No significant differences were observed in the phenotypes of *D. immitis* infection between OB and OB-wMel. The mosquito survival rate at 13 dpi was 52.1% (25/48) in OB and 63.6% (28/44) in OB-wMel (Fig 2B). The prevalence rate of L3 was 95.8% (23/24) in OB and 100% (27/27) in OB-wMel (Fig 2C). The number of L3 was 17.8 in OB and 20.1 in OB-wMel (Fig 2D). We also investigated the effects of wMel infection on the *D. immitis* resistance phenotype. The prevalence of L3 was 100% (30/30) in OB, 0% (0/30) in IB12, and 5.7% (2/35) in OB-wMel × IB12 (Fig 2E). The number of L3 in IB12 and OB-wMel × IB12 were 0 and 0.06, respectively, and they maintained the resistant phenotype. The L3 number in IB12 and OB-wMel × IB12 was significantly lower than that in OB (18.5) (Fig 2F).

**Fig 2.**
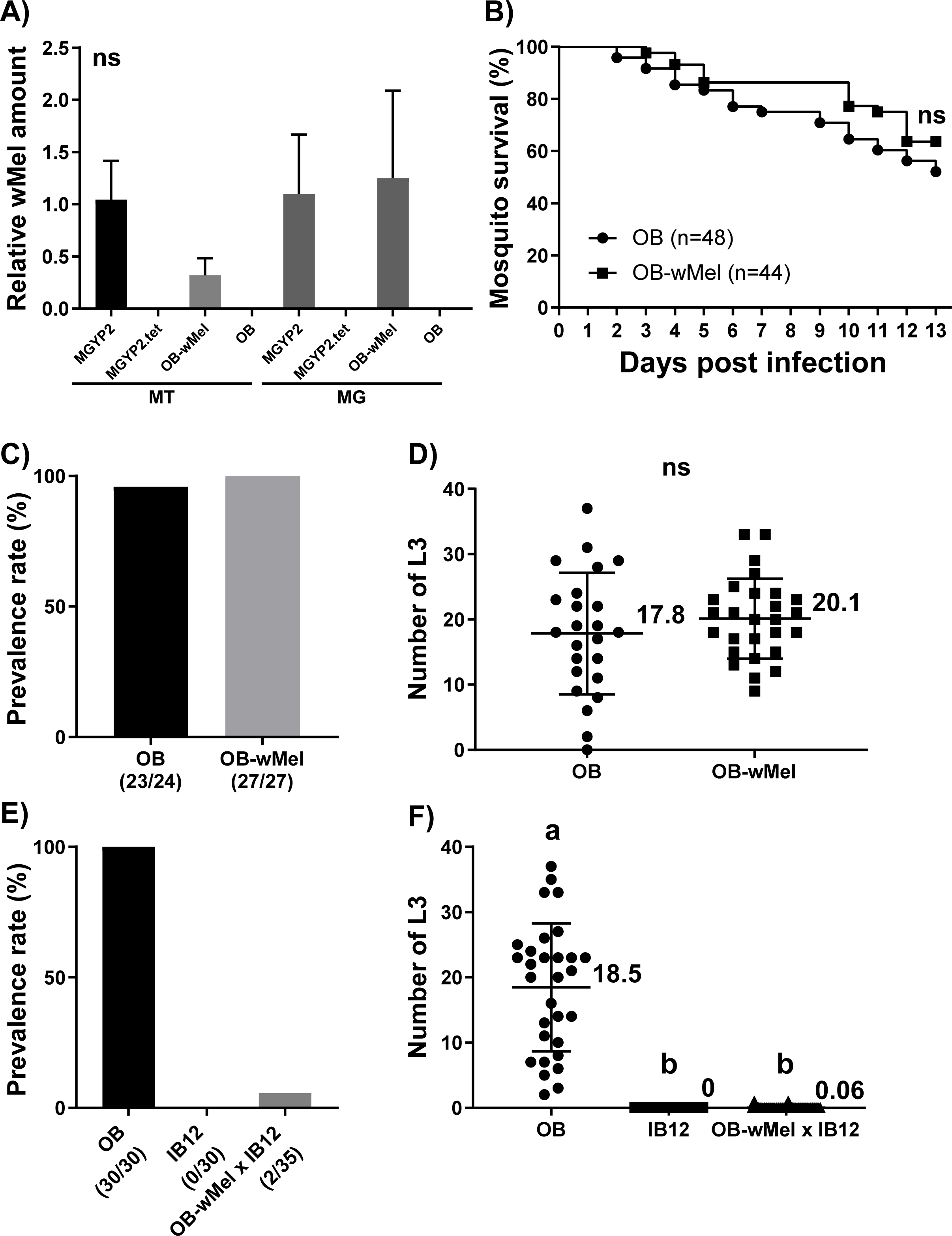
wMel had no effect on the *D. immitis* infection phenotype in both susceptible and resistant strains. *D. immitis* infection phenotypes in susceptible OB and resistant IB12 strains were compared with and without wMel. (A) Relative wMel amount in the MT and MG in MGYP2.tet, MGYP2, OB, and OB-wMel. Vertical transmission of wMel in the backcross OB-wMel was confirmed by qRT-PCR. OB and MGYP2.tet were used as negative controls. MGYP2 was used for positive control. (B) Mosquito survival up to 13 dpi in OB and OB-wMel. (C) Prevalence rate of L3 on 13 dpi in OB and OB-wMel. (D) Number of L3 on 13 dpi in OB and OB-wMel. (E) Prevalence rate of L3 on 13 dpi in OB, IB12, and OB-wMel x IB12. (F) Number of L3 on 13 dpi in OB, IB12, and OB-wMel x IB12. Experimental sample size in the analysis of mosquito survival and number of L3-positive/total as a measure of the prevalence rate are shown for each strain. Data for the relative wMel amount and number of L3 are shown as the mean value with SD. The wMel amount, mosquito survival rate, and number of L3 were statistically analyzed. Different letters indicate significant differences (*p* < 0.05). “ns” indicates no significant difference.

L3 from OB-wMel showed normal morphology similar to OB (Fig 3A). The *in vitro* L3 molting assay showed no significant difference between OB and OB-wMel (Fig 3B). The L3 molting rate on day 7 was 86.7% (26/30) in OB and 80% (24/30) in OB-wMel. These results indicate that wMel infection had no positive or negative effects on the vectorial capacity of *Ae. aegypti* for *D. immitis*.

**Fig 3.**
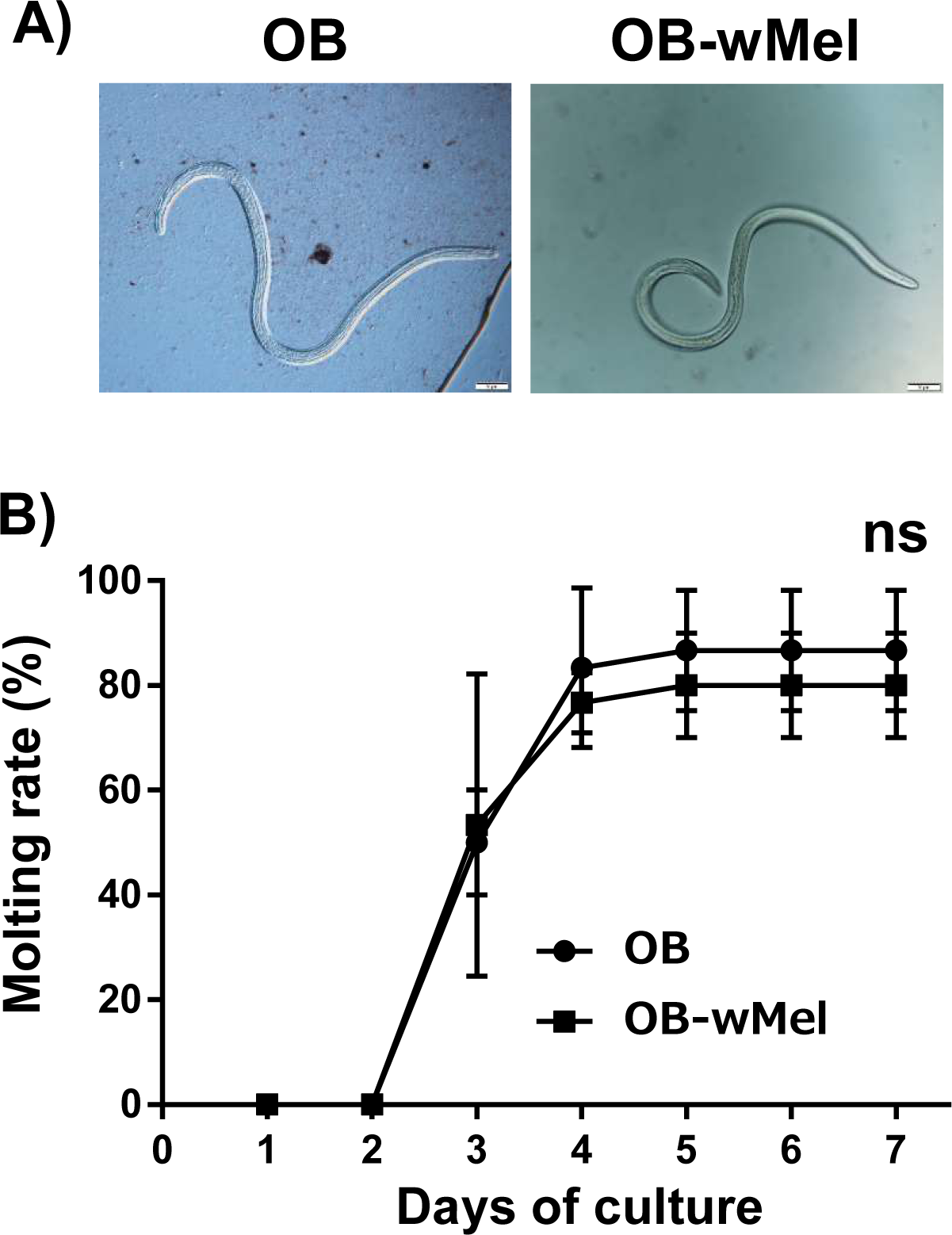
Normal molting *in vitro* in OB-wMel-derived L3. The *in vitro* molting rates of L3 were compared between the OB and OB-wMel groups. (A) Photographs indicate the normal L3 from OB and OB-wMel. A scale bar of 50 μm is shown at the lower right of the photograph. (B) Graph showing the L3 molting rate for up to 7 d of culture. The molting rate on each cultured day is shown as a mean value with SD. “ns” indicates no significant difference.

### Mosquito strain-specific gene expression patterns during co-infection with wMel and *D. immitits*

The MA plot showed that the number of significant DEGs between strains (MGYP2.tet and MGYP2, OB and OB-wMel, MGYP2.tet and OB, and MGYP2 and OB-wMel) was 190 between MGYP2.tet and MGYP2 (Fig 4A), 1075 between OB and OB-wMel (Fig 4B), 282 between MGYP2.tet and OB (Fig 4C), and 0 between MGYP2 and OB-wMel (Fig 4D). GO analysis revealed the categories of the gene functions corresponding to the DEGs detected in each strain (Table 1). The DEGs upregulated in MGYP2.tet in comparison with MGYP2 were categorized under coiled coil. The DEGs upregulated in MGYP2 in comparison with MGYP2.tet, and those upregulated in MGYP2.tet in comparison with OB were categorized as intracellular (GO:0005622). The DEGs upregulated in OB in comparison with MGYP2.tet included those involved in serine-type endopeptidase inhibitor activity (GO: 0004867) and the extracellular space (GO:0005615). The DEGs upregulated in OB in comparison with OB-wMel were categorized under coiled coil, alpha crystallin/Hsp20 domain (IPR002068), alpha crystallin/heat shock protein (IPR001436), HSP20-like chaperone (IPR008978), small heat shock protein, beta-1 type (PIRSF036514), and disulfide bonds. The heatmaps show the gene expression patterns related to the immune response in each strain. The genes upregulated by Toll activation, *Wolbachia* infection, and *B. malayi* infection tended to differ more between the MGYP2 and OB strains than between strains with and without wMel infection (Fig 5A-C).

**Fig 4.**
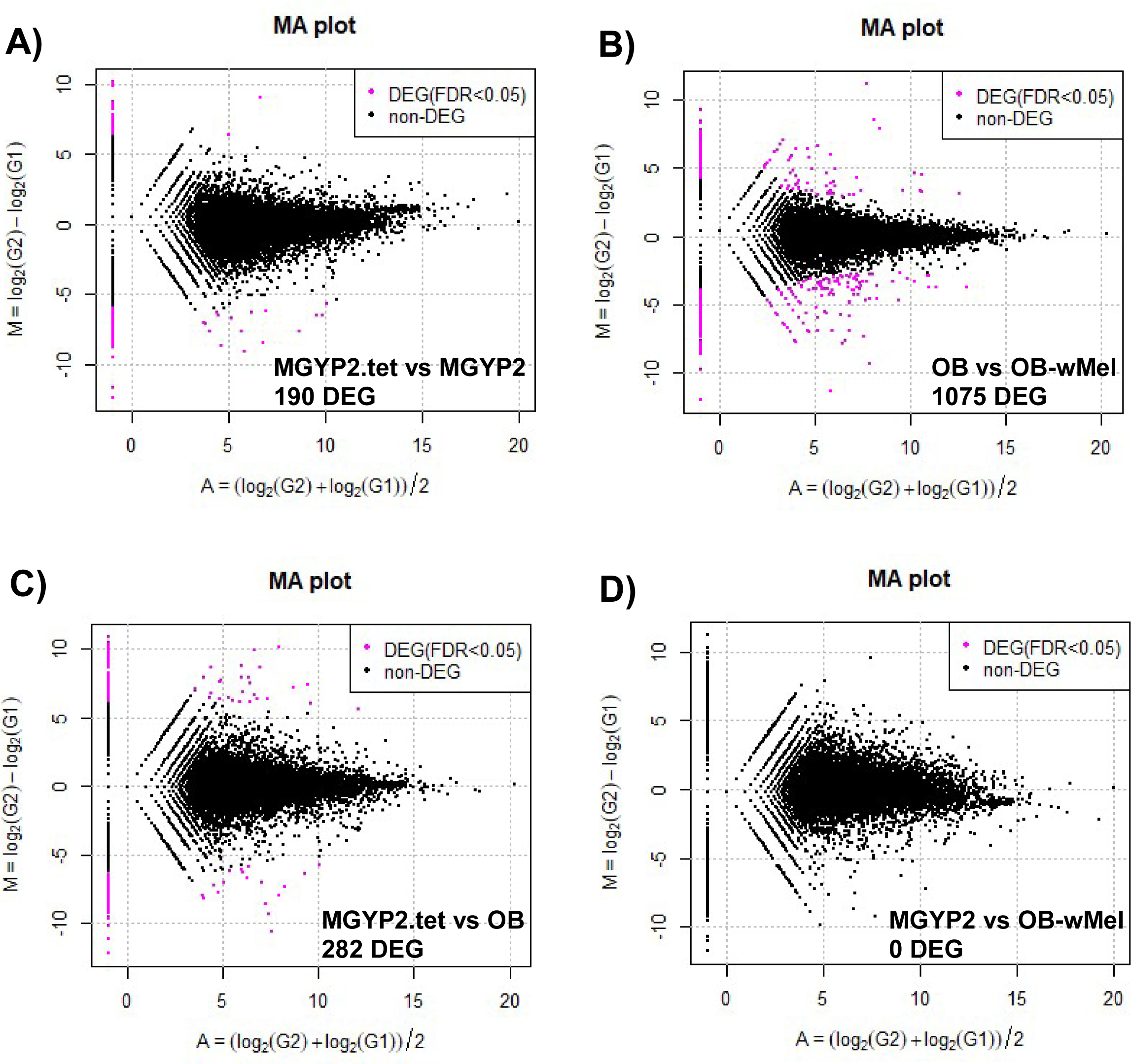
Analysis of DEGs between strains with and without wMel. An MA plot was generated to detect DEGs with false discovery rate (FDR) < 0.05, using RNA-seq data and R software. Pink dots indicate (A) 190 DEGs between MGYP2.tet and MGYP2, (B) 1075 DEGs between OB and OB-wMel, (C) 282 DEGs between MGYP2.tet and OB, and (D) 0 DEGs between MGYP2 and OB-wMel.

**Table 1.**
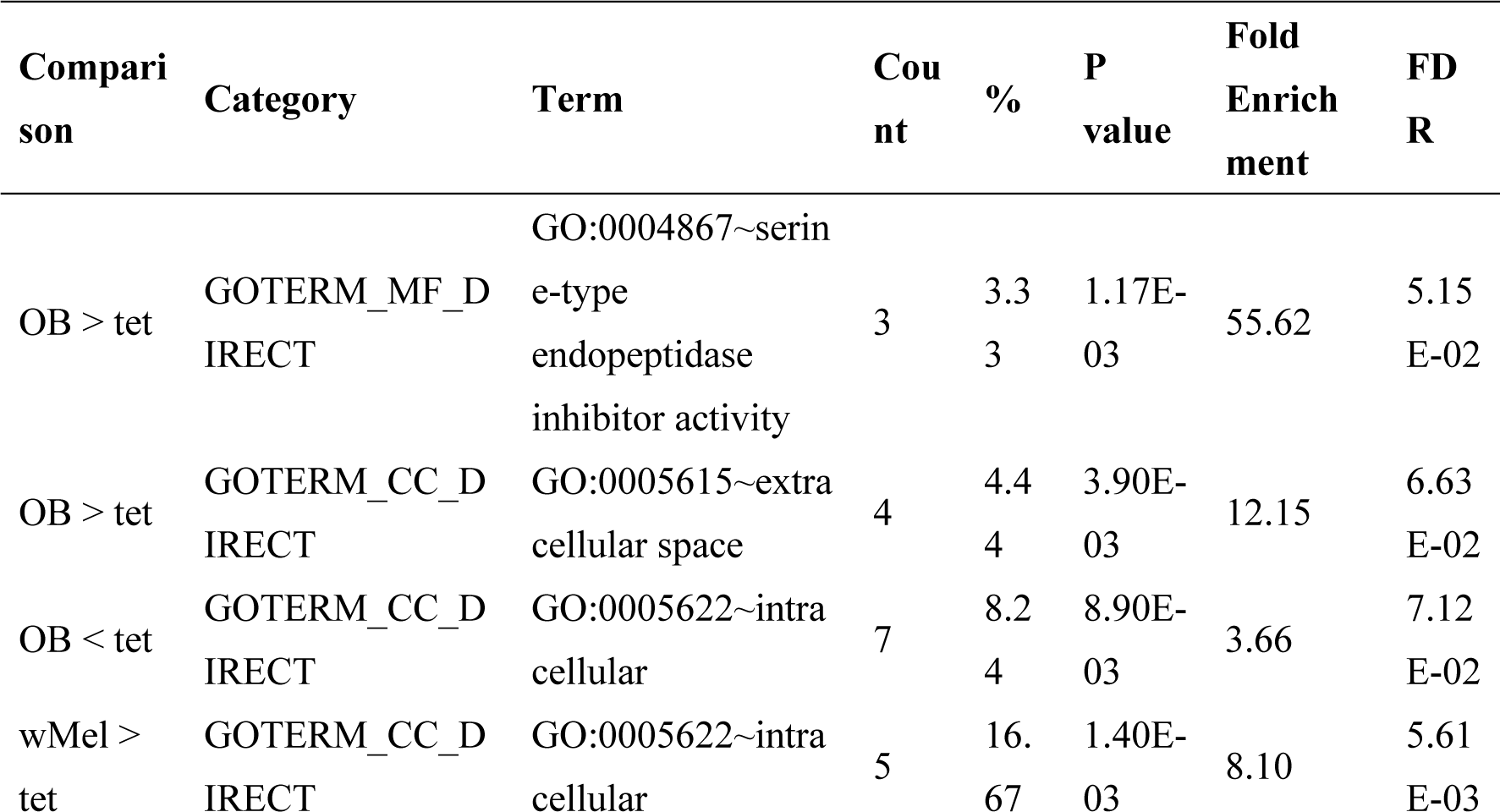
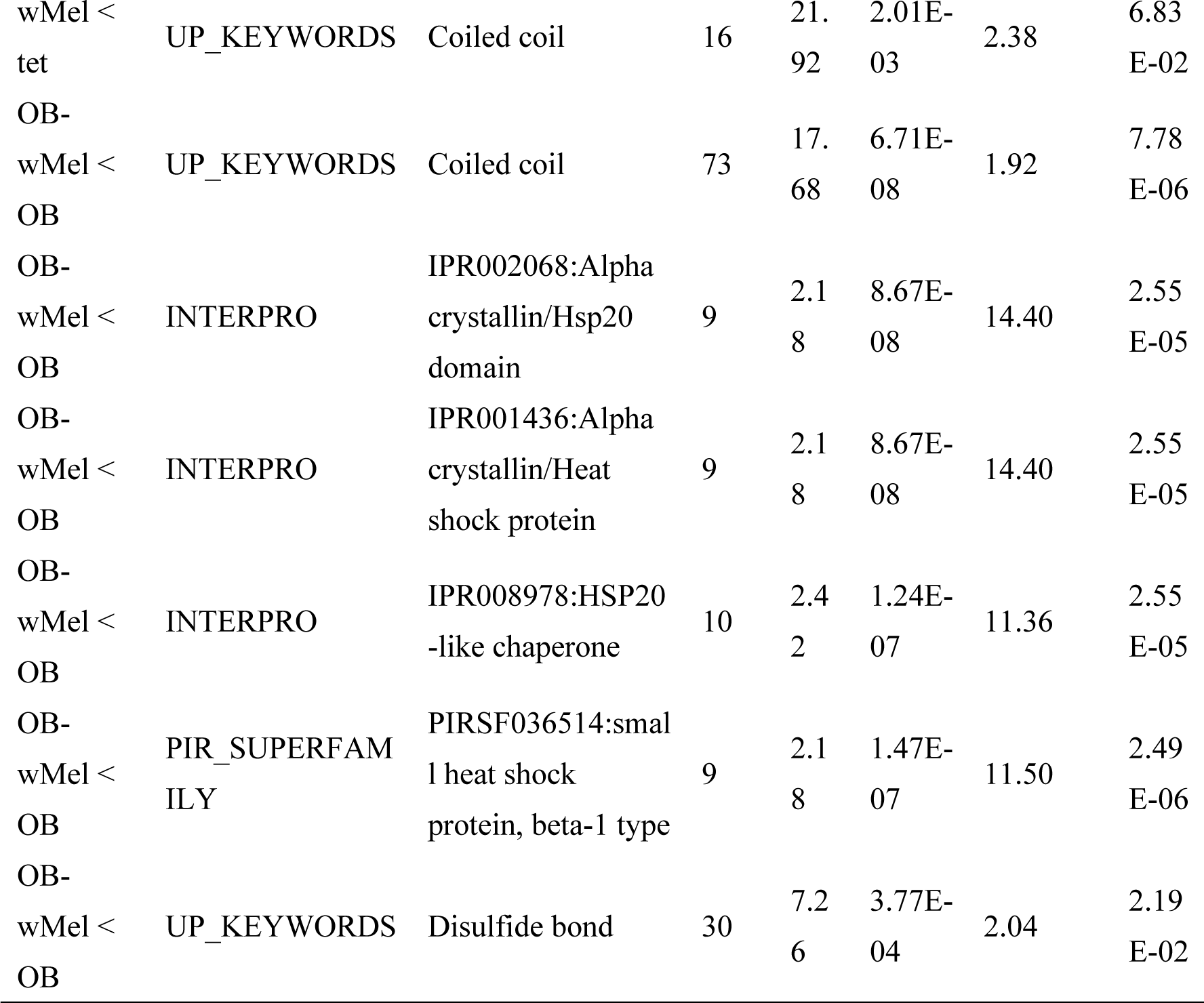
Gene Ontology enrichment analysis of DEGs between each strain by DAVID.

**Fig 5.**
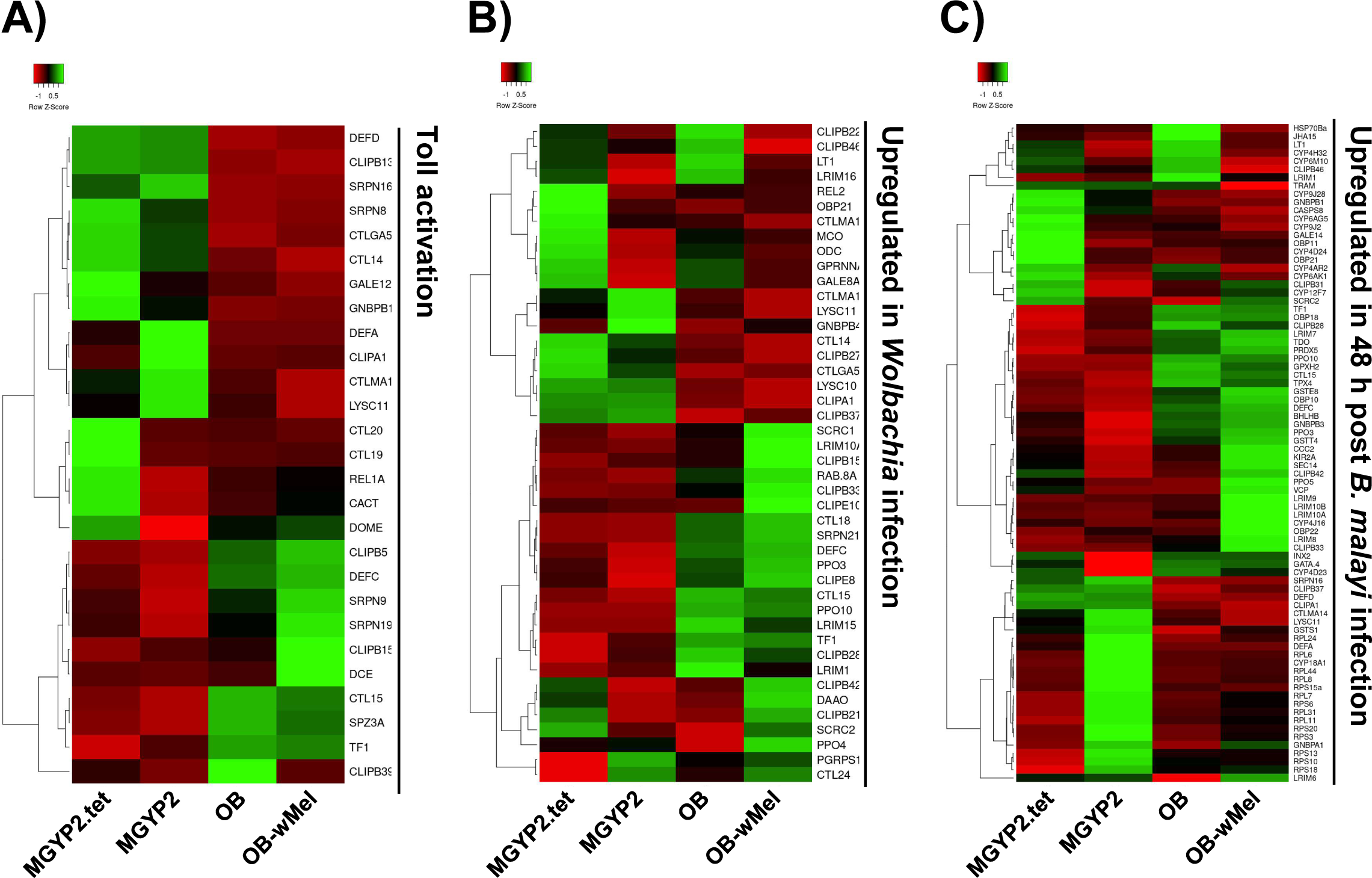
Heatmap of immune response-related gene expression showing the differences between OB and MGYP2. Heatmaps were generated for the genes that have been previously reported to be upregulated in the immune response in *Ae. aegypti*. Each figure indicates the expression tendency of genes related to (A) Toll activation, (B) *Wolbachia* infection, and (C) *B. malayi* infection in MGYP2.tet, MGYP2, OB, and OB-wMel. Darker red colors indicate lower values and darker green colors indicate higher values in the heatmap.

### Influence of *D. immitis* infection on the amount of wMel in mosquitoes

The amount of wMel DNA in the SG and MT of non-blood feeding mosquitoes on day 14 was significantly lower than the corresponding values obtained pre-blood feeding on day 0 (Fig 6A and B). No significant difference was observed between the relative amounts of wMel DNA in OB-wMel mosquitoes infected with *D. immitis* and those that were not (Fig 6A-C). The amount of wMel did not change during the course of *D. immitits* infection from mf to L3. These results suggest that *D. immitits* infection had no adverse effect on mosquito-borne disease control using *Wolbachia*, at least not by reducing the amount of *Wolbachia* in the mosquitoes.

**Fig 6.**
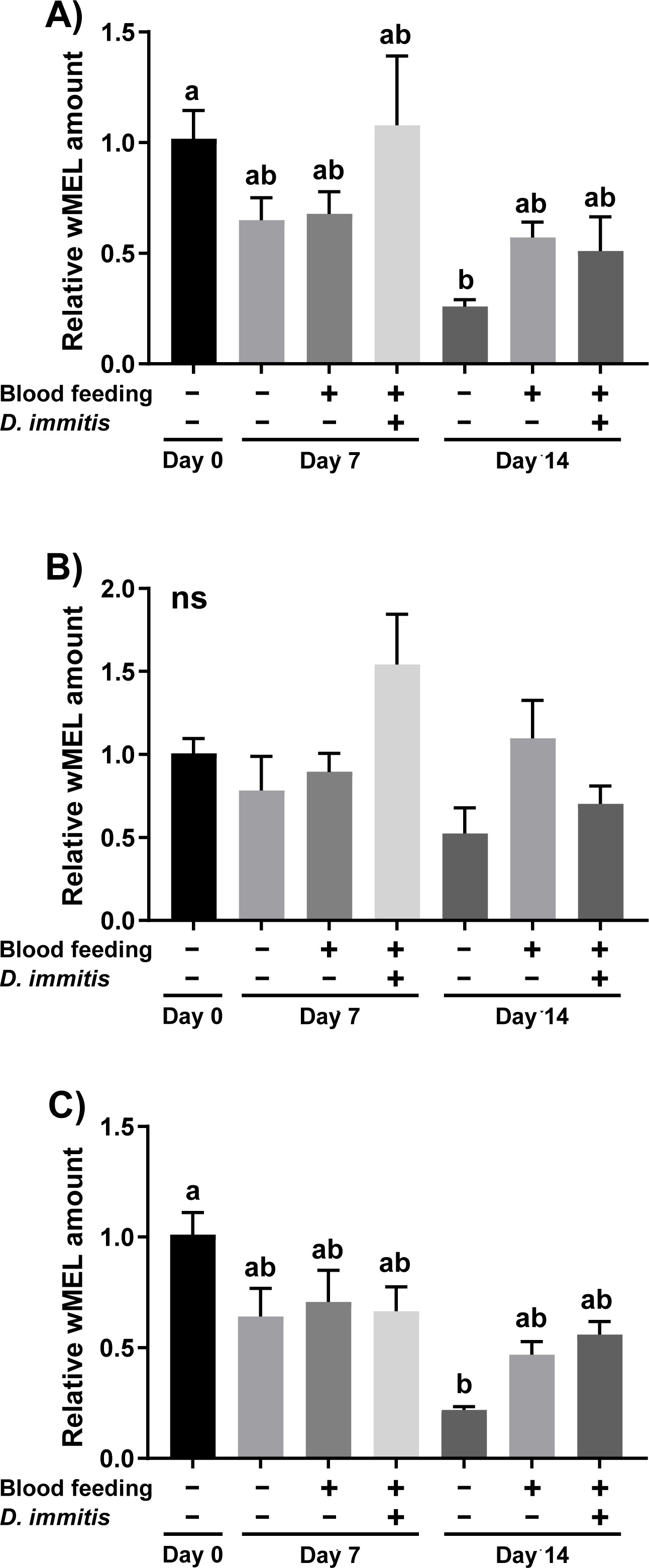
*D. immitis* infection had no effect on the amount of wMel in mosquitoes. The relative amounts of wMel in OB-wMel infected with and without *D. immitis* were analyzed by qRT-PCR. Graphs show the wMel amount in (A) SG, (B) MG, and (C) MT between pre-blood feeding on day 0 (blood feeding: -, *D. immitis*: -), non-blood feeding on days 7 and 14 (blood feeding: -, *D. immitis*: -), non-infected blood feeding on days 7 and 14 (blood feeding: +, *D. immitis*: -), and *D. immitis*-infected blood feeding on days 7 and 14 (blood feeding: +, *D. immitis*: +). All data are shown as mean with SD. The amount of wMel in each treatment group was statistically analyzed. Different letters indicate significant differences (*p* < 0.05). “ns” indicates no significant difference.

## Discussion

This study showed that wMel infection did not adversely affect the *D. immitis* infection phenotype, including mosquito survival, prevalence rate, L3 number, and L3 molting rate, in *Ae. aegypti.* Therefore, the dengue control method using *Wolbachia* is safe because it does not spread *D. immitis* infection. The use of *Wolbachia* as a dengue control strategy has been widely studied; however, to the best of our knowledge, this is the first study to investigate the effect of *Wolbachia* transfection on the vectorial capacity of *Ae*. *aegypti* for *D. immitis*. We also found no adverse effects of *D. immitis* infection on the amount of *Wolbachia* in *Ae. aegypti*. Therefore, our results suggest that a *Wolbachia*-based control strategy against dengue may be effective in endemic areas where *D. immitis* infection is possible.

The current understanding of the phenotype in the MGYP2 strain is limited because it shows a mixture of susceptibility and resistance to *D. immitis* infection [42]. To address this issue, we successfully established a highly susceptible backcross strain, OB-wMel. OB-wMel maintained the susceptibility phenotype of the original OB strain. The crossing of OB-wMel with the refractory strain IB12 resulted in a resistant phenotype. These findings suggest that wMel has no positive or negative effect on the vectorial capacity for *D. immitis* in *Ae. aegyptI*. Similar results were reported for *B. pahangi* in *Ae. polynesiensis* [32] and *Ae. pseudoscutellaris* [33]. However, no previous studies on the phenotypic analysis of *D. immitis* in *Wolbachia*-infected *Ae. aegypti* have been reported to date, and the wMel infection in *Aedes* may not promote an active host response to *D. immitis* infection. Therefore, further studies are required to support our findings. The amount of wMel in the mosquitoes did not change during *D. immitis* infection, suggesting that *D. immitis* infection does not interfere with the control of mosquito-borne diseases in *Wolbachia*-infected mosquitoes. Although *Wolbachia*-infected mosquitoes are not expected to be effective in controlling filariasis, they have no adverse effects on *D. immitis* infection, and the control of dengue fever using *Wolbachia* can be recommended as a safe method.

The effect of *Wolbachia* on host response in mosquitoes differs by strain. *Ae. aegypti* infected with the virulent strain wMelPop have a shorter lifespan than those infected with wMel [43–45]. Thus, stimulation of the immune system is stronger in wMelPop than in wMel [31, 40, 44, 46]. In the present study, the expression of immune response-related genes after *D. immitis* infection was not significantly different between wMel-infected and non-infected mosquitoes. However, wMel infection showed no effect on the vectorial capacity of *Ae. aegypti* for *D. immitis* in this study, whereas wMelPop infection suppressed the vectorial capacity of *Ae. aegypti* for *B. pahangi* in a previous study [31]. *P. berghei* oocyst levels in *An*. *gambiae* were suppressed by wMelPop and the opposite effect was observed for wAlbB [21]. Thus, mosquitoes infected with wMelPop may activate their immune system more than those infected with wMel, suppressing their vectorial capacity for *D. immitis*.

The suppressive effect of *Wolbachia* on the vectorial capacity of mosquitoes also depends on the experimental conditions, such as the combination patterns of *Wolbachia* strain, mosquito species, and pathogen types [5]. Therefore, investigating the immune response of mosquitoes against target pathogens is essential when using *Wolbachia*. The *Wolbachia* surface protein can trigger innate immune activation in insects and mammals [47, 48]. *Ae. aegypti* infected with the transgenic symbiotic bacterium *Asaia* expressing the *Wolbachia* surface protein showed reduced vectorial capacity for *D. immitis* [34]. The main components of the immune system of mosquitoes are the Toll and Imd pathways, antimicrobial peptides, and melanization. The Toll pathway and antimicrobial peptides are essential for defense against dengue virus infection [49, 50], while melanization is important for defense against parasitic infections. Recently, Toll activation has been reported to block the development of *D. immitis* and *B. malayi* in *Ae. aegypti* [37]. Confirming the activation of effective immune response pathways against the pathogen in *Wolbachia*-infected mosquitoes is important for ensuring appropriate practical applications in mosquito-borne disease control. Transfection of suitable *Wolbachia* strains or the above transgenic bacteria into mosquitoes has the potential to suppress the vectorial capacity of several pathogens, including *D. immitis*.

The present study showed that wMel-infected *Ae. aegypti* maintained the phenotype of *D. immitis* infection. However, the selection of a suitable *Wolbachia* strain to enhance the immune system in mosquitoes can be applied to mosquito-borne diseases other than dengue [5]. Our results also show that *D. immitis* infection did not affect the amount of wMel in *Ae*. *aegypti*. The endemic areas of dirofilariasis and dengue fever are common, mainly in Southeast Asia and South America, and are expected to expand in the future due to global warming [24, 51]. Therefore, our results suggest that control of dengue using *Wolbachia*-transinfected mosquitoes is an effective approach that poses no risk of spreading of filariasis or interfering with dengue control in common endemic areas of dengue and filariasis. Previous studies have also found that the effectiveness of *Wolbachia* in suppressing vectorial capacity in *Ae*. *aegypti* was not affected by co-infection with dengue and Zika viruses [52]. Future studies should investigate the effects of co-infection with dengue virus and other pathogens, including filaria, on the vectorial capacity of both pathogens in *Wolbachia*-infected mosquitoes.

## Acknowledgments

We are deeply grateful to Dr. Scott O’Neill, Professor at Monash University, for providing the MGYP2 strain. We also thank Dr. Koji Kadota, Associate Professor at Tokyo University, for the advice on the RNA-seq data analysis. We thank the students at OUAVM for their cooperation in maintaining the wMel-infected mosquitoes. The following reagent was obtained through BEI Resources, NIAID, NIH: *Aedes aegypti*, Strain LVP-IB12, Eggs, MRA-735, contributed by David W. Severson. This study weas supported by the JSPS KAKENHI (grant numbers 22H02510, 22K19235 and 19KK0175 to SF).

